# Augmentation of Virtual Endoscopic Images with Intra-operative Data using Content-Nets

**DOI:** 10.1101/681825

**Authors:** A. Esteban-Lansaque, Carles Sanchez, Agnes Borras, Debora Gil

## Abstract

Medical imaging applications are challenging for machine learning and computer vision methods, in general, for two main reasons: it is difficult to generate reliable ground truth and databases are usually too small in size for training state of the art methods. Virtual images obtained from computer simulations could be used to train classifiers and validate image processing methods if their appearances were comparable (in texture and color) to the actual appearance of intra-operative medical images. Recent works focus on style transfer to generate artistic images by combining the content of an image and the style of another one. A main challenge is the generation of pairs with similar content ensuring preservation of anatomical features, especially across multi-modal data. This paper presents a deep-learning approach to content-preserving style transfer of intra-operative medical data for realistic virtual endoscopy. We propose a multi-objective optimization strategy for Generative Adversarial Networks (GANs) to obtain content-matching pairs that are blended using a siamese u-net architecture (called Content-net) that uses a measure of the content of activations to modulate skip connections. Our approach has been applied to transfer the appearance of bronchoscopic intra-operative videos to virtual bronchoscopies. Experiments assess images in terms of, both, content and appearance and show that our simulated data can substitute intra-operative videos for the design and training of image processing methods.

## Introduction

State of the art methods in computer vision need huge amounts of data with unambiguous annotations for their training. In the context of medical imaging this is, in general, a very difficult task due to limited access to clinical data, the time required for manual annotations and variability across experts. The particular field of intervention guiding has the extra difficulty of intra-operative recordings probably requiring the alteration of standard protocols. These facts have encourage the development of computational methods for the simulation of realistic medical imaging data from the available scans. In this context, virtual endoscopy could be useful to train intervention support methods.

In order to obtain realistic data useful for data augmentation and validation of machine learning and image processing methods, simulations should resemble intra-operative recordings. We consider that style transfer could be used to endow virtual endoscopic images with the content and texture of intra-operative videos using modern techniques for artistic style transfer.

The basic idea of style transfer [1, 2] is to transform input images appearance according to one or more style images while preserving structure and content of input images. Recent works [3, 4] have shown the power of Generative Adversarial Networks (GANs) and Convolutional Neural Networks (CNNs) in general for artistic style transfer.

The main difference between artistic style transfer and realistic simulations of endoscopic procedures is that in the latter, stylized images should preserve the structure of simulated data. This follows from the fact that style structure encodes the anatomical content of the image and, thus, it should be preserved.

This work addresses the generation of realistic endoscopic images using intra-operative video data to augment the appearance of virtual endoscopy. We present a two-stage method based on CNNs that maps virtual images to the intra-operative domain preserving their anatomical content.

### Related work

State-of-art techniques for artistic style transfer based on CNNs use a system of two different neural networks (a generative network and a discriminative one) to obtain stylized images achieving a compromise between preservation of input image content and style images texture and appearance. The first network is an auto-encoder that generates stylized images from the input images. The output of this auto-encoder is the input of a discriminative network that classifies between style and input images to assess how much stylized images appearance matches the actual style. This general scheme has several variants concerning, especially, the kind of loss function used to train the network.

In [3, 5] the generative network minimizes a loss function that includes two terms. One term (feature reconstruction loss) penalizes that stylized images deviate in content from input images, while the second one (style reconstruction loss) measures the similarity between stylized images and style appearance. The feature reconstruction loss is given by the *L*_2_ difference between feature maps of input and stylized images. The style reconstruction loss is given by the Frobenius norm of the difference between the Gram matrices of the stylized and style images feature maps. As explained in both works, since Gram matrices encode probabilistic distribution correlations, minimizing the style reconstruction loss preserves stylistic features from the style image but does not necessarily preserve its spatial structure (as illustrated in Fig 1(b)).

**Fig 1.**
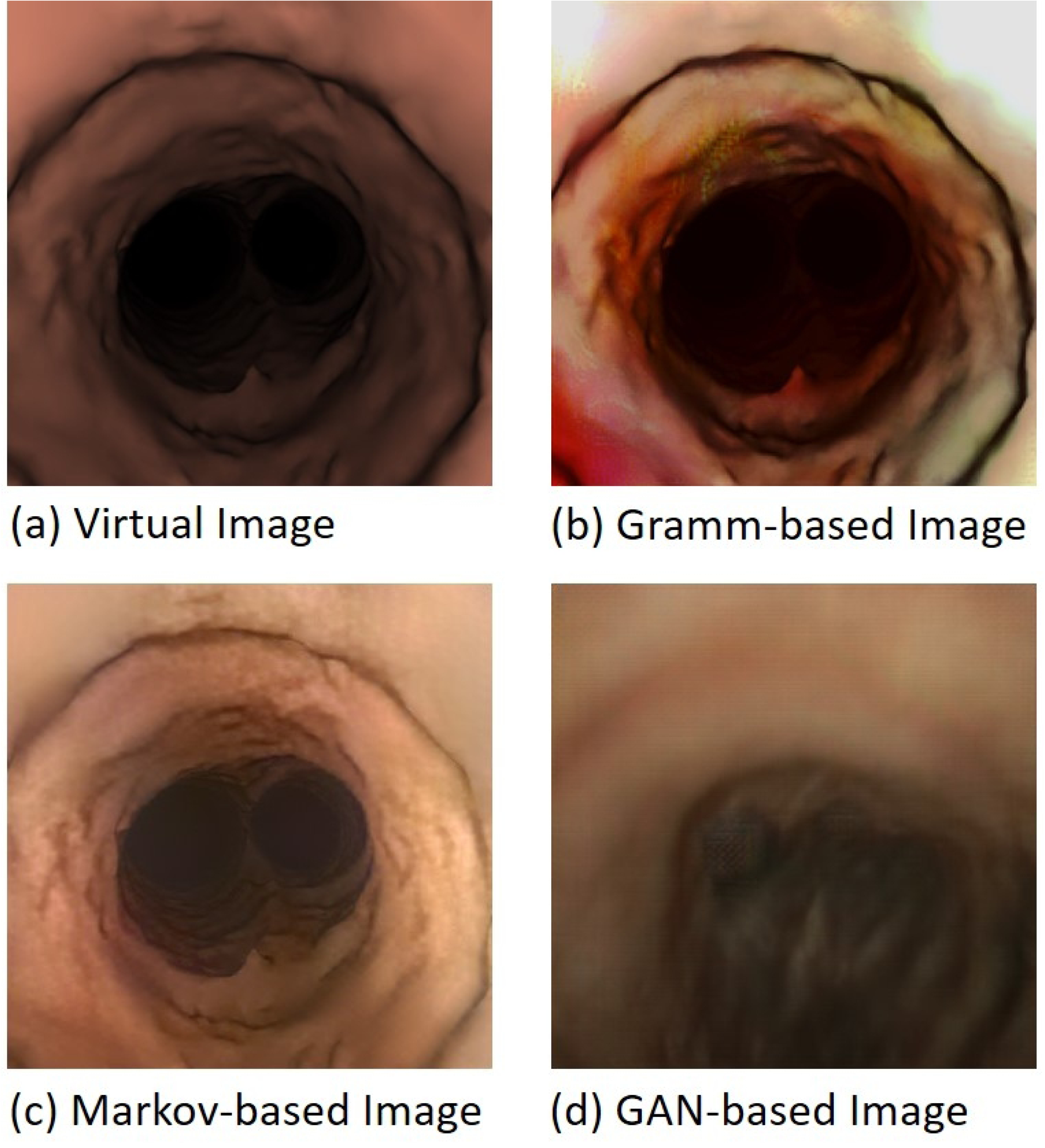
Virtual image stylized with different state-of-art methods.

Other approaches [6] are based on deep markovian models and transform images locally instead of globally. To do so, feature maps are split into patches which are the input of the classifier that discriminates between real and virtual appearances. Like [3, 5] the loss function also includes a content regularization term to preserve the spatial structure of images. However, the fact that style is locally transferred also leads to a loss of anatomical content in style images (Fig 1(c)).

New approaches such as [4] use Generative Adversarial Networks (GANs) in order to transform images from one domain *A* (like virtual simulations) into a domain *B* (like interventional videos). The novelty of [4] is that a cyclic term is added in order to make the domain transfer bijective (*A*→ *B* → *A* and *B* → *A* → *B*). Although, the method also adds a regularization term to preserve the spatial structure of the stylized virtual images, content information is still lost as shown in Fig 1(d).

### Contributions

We present a two-stage algorithm for the augmentation of virtual endoscopic images using intra-operative videos based on convolutional neural networks. First, we use cycleGAN in a multi-objective optimization scheme to obtain pairs of virtual and intra-operative style images that share some content information. The content and appearance of these image pairs are blended using a siamese u-net architecture that modulates skip connections by a measure of neuron activation content. The contributions of our work are the following:

1. *A multi-objective approach to GANs*. A main challenge with GANs [7] is the selection of the epochs most suitable for a given problem. Due to the oscillating behavior of adversarial training, most of the cases this is done manually, which leads to subjective errors depending on the observer. In this work, we propose a multi-objective optimization approach based on the Pareto front [8] to select the epoch achieving the best compromise between content preservation and style transfer.
2. *A siamese u-net network (ContentNet) for blending content and appearance of image pairs*. We introduce an auto-encoder with siamese encoders and skip connections to the decoder. Skip connections are modulated by a measure of the information according to the type of information filters respond to. Modulation is used to fine tune the amount of virtual anatomical content and intra-operative appearance of final simulations. This way we can produce images with several degrees of interventional artifacts from the same pair of images obtained from multi-objective GANs.

## Materials and methods

### Augmentation of Virtual Endoscopic Images using Intra-operative Data

To obtain realistic textured endoscopic images is a very complicated task since it is usually very difficult to have exact correspondences between real and virtual images. Besides methods for unpaired mapping fail to preserve the anatomical content of the original domain.

Our strategy for intra-operative virtual endoscopy is a two-step method. In a first stage, we generate pairs of virtual and intraoperative images sharing anatomical content using GANs. In a second step, the content and appearance of such pairs are blended using a siamese u-net architecture trained to modulate the amount of content and texture that is taken from each pair.

#### Multi-Objective Generative Adversarial Networks

Given two domains *Virtual, V*, and *Real, R*, a GAN learns two (bijective) maps (*G*_*r*_, *G*_*v*_) from one domain onto the other one:

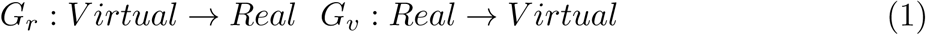

with the map composition *G*_*r*_(*G*_*v*_) and *G*_*v*_(*G*_*r*_) being the identity on each domain. Following [4], maps are given by auto-encoders trained to optimize:

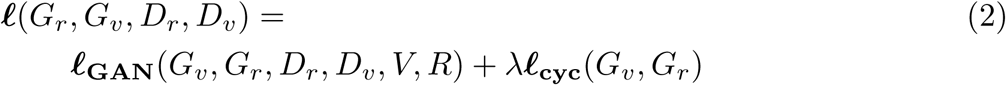

The term **ℓ**_**GAN**_ measures how good are *G*_*v*_, *G*_*r*_ transferring images from one domain to the other one, while **ℓ**_**cyc**_ is a “cycle consistency loss” introduced to force bijective mappings.

The minimization problem is given by adversarial training as:

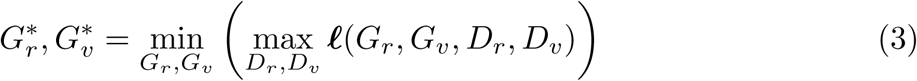

This way, 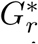 and 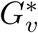 are optimized so that *G*_*r*_, *G*_*v*_ minimize (2) while the adversarial *D*_*r*_, *D*_*v*_ maximize it. These conditions (minimize while maximizing at the same time) might be in conflict when they are considered as a single optimization process. Such a conflict is prone to introduce an oscillating behavior in back propagation iterations, which might hinder the convergence of cycleGAN training and the selection of the stopping epoch.

We propose to consider separately the optimization of each of the terms in the objective function **ℓ** and pose adversarial training as the following multi-objective optimization [8] problem:

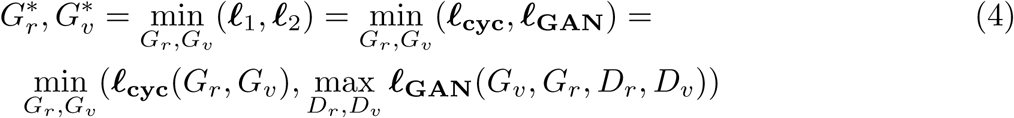

Since a multi-objective optimization problem involves the optimization of multiple objective functions, there does not exist, in general, a solution simultaneously minimizing all of them. The expected situation is to have a set of solutions that outperform in any of the objectives without degrading the other ones. These dominating solutions is called Pareto front and is defined as the set of solutions, *x*_*P*_, such that:

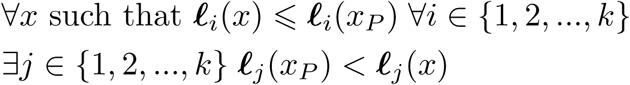

being **ℓ**_1_, …, **ℓ**_*k*_ the set of functions to be optimized.

The condition of the Pareto front can be used to select cycleGAN epoch as follows. Let 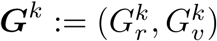 be the transformation maps at the *k*-th epoch and 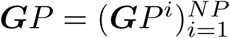 be the set of epochs belonging to the Pareto front. Such Pareto maps can be iteratively computed from the values of the objective functions as:

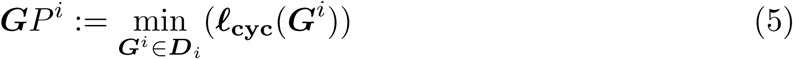

for ***D***_1_ the set of maps for all epochs and ***D***_*i*_, *i* > 2, the set of maps dominating ***G****P*^*i-*1^. In our case, ***G***^*i*^ ∈ ***D***_*i*_ if it satisfies the following conditions:

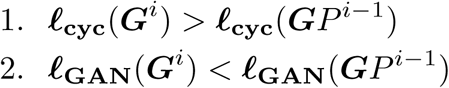

Fig 2 shows the Pareto front associated to the two-objective problem given by (4) with the dashed lines enclosing the region that includes the set of epochs that dominate a given ***G****P*^*i-*1^. The pairs of function values (**ℓ**_**cyc**_(***G****P*^*i*^), **ℓ**_**GAN**_(***G****P*^*i*^)), ***G****P*^*i*^ ∈ ***G****P*, given by the two objectives evaluated at the Pareto front are shown in solid squares joined with a solid line.

**Fig 2.**
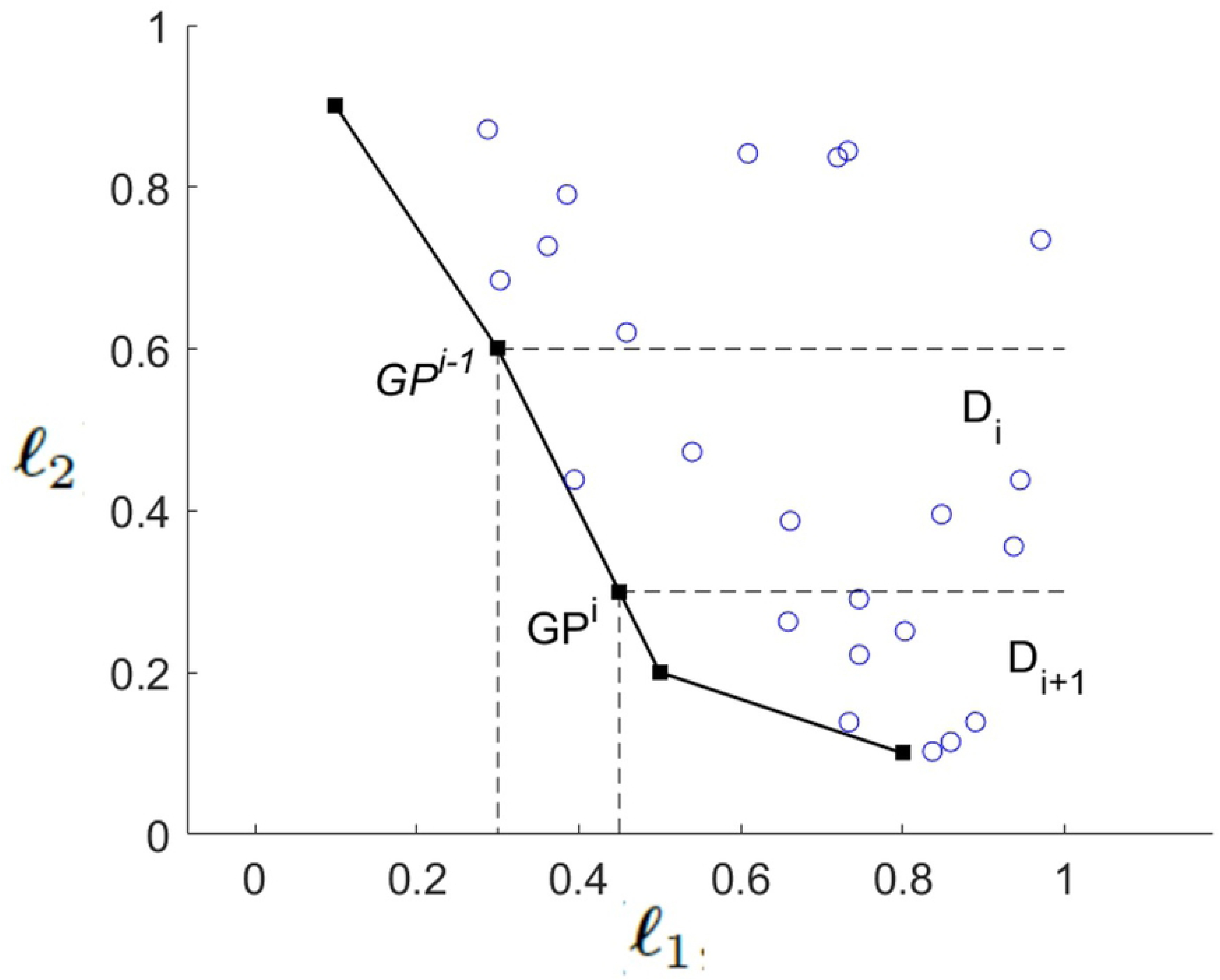
Pareto front of cycle-GAN 2-objective optimization.

The set of Pareto epochs achieve the best trade-off between the two objective functions and, thus, are equivalent from the point of view of the GAN. The epoch from the Pareto front best suited for augmentation of virtual endoscopic images is selected as the one that minimizes the *L*^2^-difference:

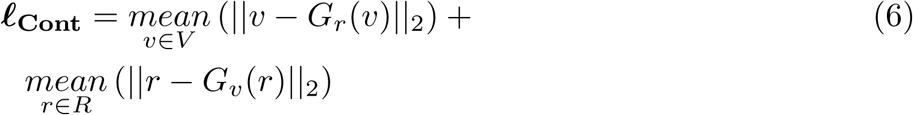

for ‖ *·* ‖_2_ the *L*^2^-norm and 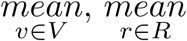 the average values for the training set of virtual, *V*, and real images, *R*. The epoch selected, 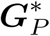 is the one in the Pareto front achieving the minimum value of **ℓ**_**cont**_:

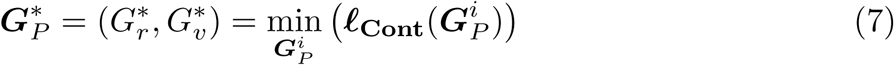

We will note the images transformed by these maps by 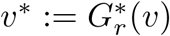 and 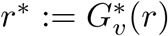.

#### ContentNet

The proposed content-net (labelled *C*) is an auto-encoder with siamese encoders (one for each image domain) that have skip connections to the decoder in order to blend content and style of image pairs. Siamese encoders follow a VGG-19 architecture. Each of them is build using VGG-19 first three convolutional blocks without the last max pooling. We decided to use only three convolutional blocks and two max poolings because all the texture that we want to transfer is practically removed after two max pooling. Content-net architecture is sketched in Fig 3.

**Fig 3.**
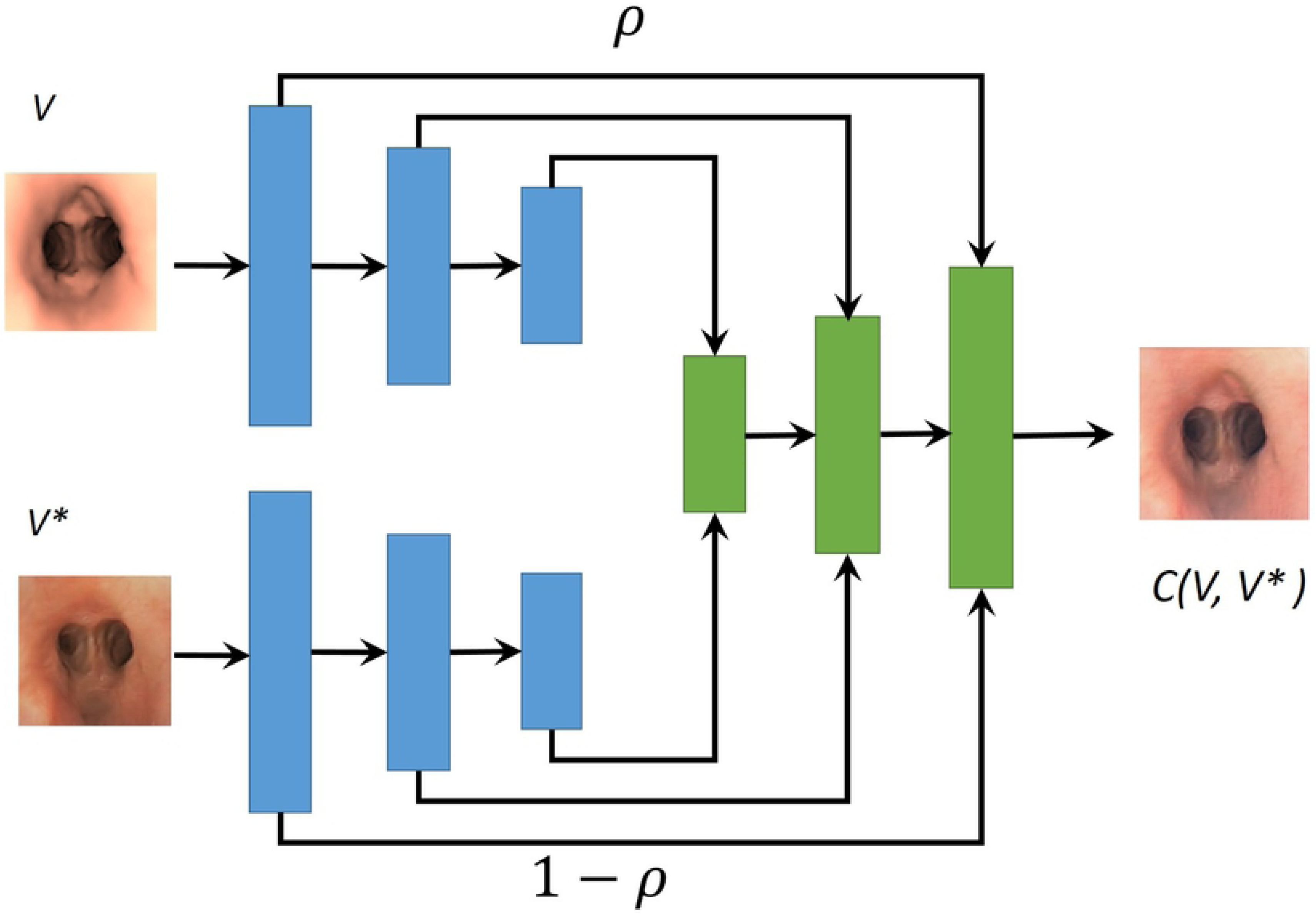
Content-net Architecture used for blending image pairs.

In order to selectively blend the content of one image with the appearance of the other one, skip-connections are weighted by a function, namely *ρ*, that quantifies the amount of content that filters respond to. Feature maps of the first siamese encoder are weighted with *ρ*, while its sister encoder are weighted with 1 − *ρ*. This way each siamese encoder contributes with either image content (first siamese encoder) or appearance (second siamese encoder).

The contrastive loss function that content-net minimizes is given by a content loss, **ℓ**_**Cont**_, defined as:

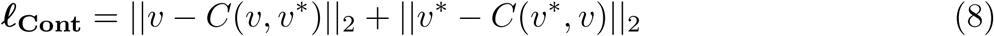

for *v* and *v*\* an image pair produced by our multi-objective cycleGAN and *C*(*v, v*\*), denoting the output of content net with *v, v*\* being inputs of, respectively, the first and second siamese encoders.

The function *ρ* weighting skip-connections is learned from a training set of intra-operative images by comparing the input image to the activation of each neuron in an encoder trained to yield the identity map. The similarity measure chosen to compare input images to its neuron activations is their mutual information [9]. Mutual information compares the correlation between random variables and is used in multimodal registration to compare images with equal content but different appearance. If **v** denotes the random variable given by input images and **a** the random variable given by each neuron activation, then their mutual information, *I*(**v, a**) is given by:

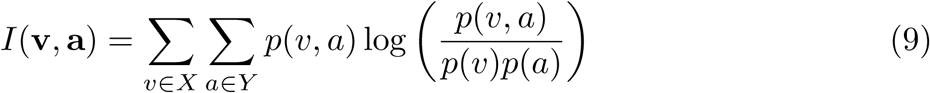

where *p*(*v, a*) is the joint probability function of **v** and **a**, and *p*(*v*) and *p*(*a*) are the marginal probability distribution functions of **v** and **a**, respectively. Mutual information measures the independence between the two random variables **v, a**. In particular, a value close to 0 indicates that input image and activations do not share information. Meanwhile, in case of sharing content, **v** and **a** would be dependent and *I* would be equal to the input image entropy.

Given an input image **v**, the mutual information evaluated for all neurons’ activations, 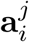, of a given layer, ℓ^*j*^, defines a strictly positive random variable:

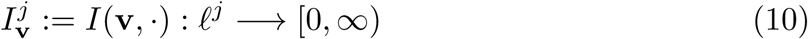

The distribution of 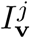 is bimodal and allows to categorize neurons’ activations into two groups sharing high and low content with the input image **v**.

The function *ρ* weighting each neuron activation is defined as the probability of a neuron sharing high content with input images and is computed as the percentage of times each activation belongs to the high content class for a set of training images:

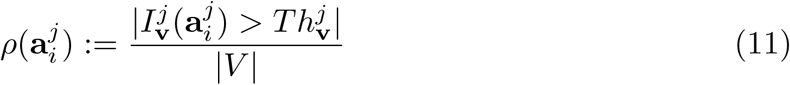

for *V* a training set, | · | indicating the length of a set and 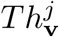 the threshold value splitting 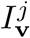 populations.

### Experiments

Our method has been used to simulate realistic bronchoscopic interventions. Virtual bronchoscopies defining the *V irtual* domain were generated using [10] from CT scans acquired with an Aquilion ONE (Toshiba Medical Systems, Otawara, Japan). Scans were selected from 10 patients in the CPAP study [11] conducted at Hospital Bellvitge (Barcelona, Spain). For each scan, we simulated 4 paths, one for each main lung lobule (left and right upper lobes; left and right lower lobes). Intra-operative videos defining the *Real* domain were acquired also at Hospital Bellvitge (Barcelona, Spain) during biopsy interventions using an Olympus Exera III HD Ultrathin videobronchoscope (6 videos) and CAO diagnostic procedures using a Olympus Exera III HD Therapeutic videobronchoscope (4 videos).

Methods were trained on 4 CT anatomies (16 virtual bronchoscopies) and video recordings from 3 ultrathin explorations and 2 diagnostic procedures. The remaining data were left for validation and testing. The multi-objective cycleGAN was trained from scratch using the whole training set. After 200 epochs, our multi-objective approach selected epoch 50 as the one achieving the best compromise between intr-operative appearance and preservation of virtual anatomical content. Content-net was fine-tuned on the set of intra-operative recordings from an auto-encoder trained to yield the identity map on the *Real* domain. The weighting function *ρ* was also learned from the latter auto-encoder.

## Results and Discussion

We evaluated the quality of the enhanced virtual images in terms of intra-operative appearance and preservation of each patient’s anatomy acquired by CT scans. For comparison purposes, simulations were also enhanced using the 200th epoch network, labelled GAN200, and the network achieving the least value of the cost (2), labelled GANLeast. This last network corresponded to the 21st epoch of cycleGAN training.

### Intra-operative Appearance

For the first experiment, we trained a network, *D*_*r*_, to discriminate between real and virtual images. This network was evaluated on the test sets of real images and virtual images enhanced using the proposed method, GAN200, and GANLeast.

For each enhancing method, *D*_*r*_ values were compared to the ones obtained by the test real images to assess their similarity in appearance. Main analysis was performed using a Student T-test for unpaired data. We computed p-values and 95% confidence interval (CI) for the difference. A p-value < 0.05 was considered statistically significant.

Table 1 summarizes statistics for the comparison of appearances. For each enhancing method, we report p-values, 95% CIs of the difference with real image and descriptive statistics (mean and standard deviation (SD)) of *D*_*r*_ values. According to T-tests, none of the methods has an appearance significantly different from intra-operative videos (p-value> 0.05) with all discrimination values *D*_*r*_ in similar ranges comparable to the values achieved by the test set of real images (mean=0.90 and SD=0.11).

**Table 1.** *D*_*r*_ Statistics for Assessment of Intra-operative Appearance.

| Method | Descriptive |  | T-test |  |
| --- | --- | --- | --- | --- |
|  | mean | SD | p-val | 95% CI |
| ContentNet | 0.88 | 0.11 | 0.305 | (-0.023,0.073) |
| GAN200 | 0.91 | 0.08 | 0.576 | (-0.057,0.032) |
| GANLeast | 0.91 | 0.09 | 0.438 | (-0.065,0.028) |

Fig 4 shows representative images of virtual bronchoscopies enhanced using our method (Fig 4 (a)), the 200th epoch network (Fig 4(b)) and the least cost one (Fig 4(c)). For each case, we show two consecutive frames of the enhanced virtual sequence which should be very similar in appearance and content. GANLeast images have sudden dark artifacts, while GAN200 yields highly unstable images that do not always match the original anatomy. ContentNet provides a stable appearance in images which are the most consistent with the original anatomical content of virtual images.

**Fig 4.**
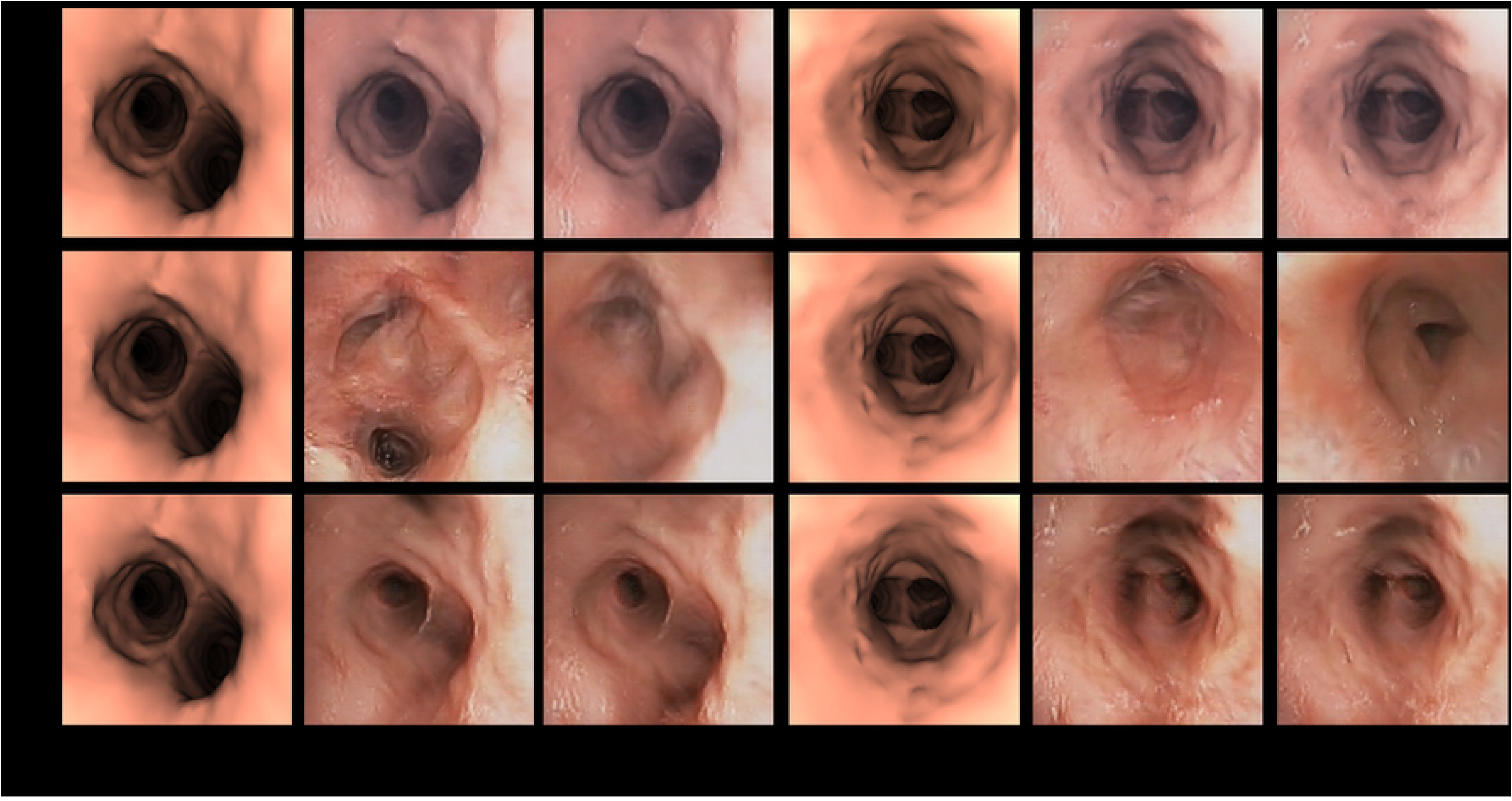
Virtual bronchoscopy enhanced using ContentNet, (a), 200th epoch network, (b) and the least cost one, (c).

### Anatomical Content

For the second experiment, we applied the lumen center detector [12] to ContentNet, GAN200, GANLeast and the non-enhanced original virtual images to verify that original lumen position and structure is preserved. The center detector was applied using two different sets of parameters, one learned on interventional videos and the other one learned on simulated bronchoscopies. Interventional parameters were used on enhanced images, while simulation parameters were applied to original virtual images. Like [12], detections were plot on original virtual images and shown to 2 independent observers for the identification of false detections and missed centres. Ground truth was produced by intersecting the experts’ annotations and used to compute precision and recall.

Scores obtained for ContentNet, GAN200, GANLeast were compared to the ones obtained for virtual non-enhanced images using a Student T-test for paired data. As in the first experiment, we computed p-values and 95% confidence intervals and a p-value < 0.05 was considered statistically significant.

Table 2 and 3 summarize statistics for Prec and Rec reported as in Table 1. ContentNet outperforms the two cycleGAN, both, in terms of precision and recall.

**Table 2.** Precision Statistics for Assessment of Anatomical Content.

| Method | Descriptive |  | T-test |  |
| --- | --- | --- | --- | --- |
|  | mean | SD | p-val | 95% CI |
| ContentNet | 0.94 | 0.09 | 0.025 | (-0.1150, -0.0082) |
| GAN200 | 0.89 | 0.15 | 0.695 | (-0.0719, 0.0487) |
| GANLeast | 0.84 | 0.19 | 0.265 | (-0.0332, 0.1148) |

**Table 3.** Recall Statistics for Assessment of Anatomical Content.

| Method | Descriptive |  | T-test |  |
| --- | --- | --- | --- | --- |
|  | mean | SD | p-val | 95% CI |
| ContentNet | 0.90 | 0.14 | 0.939 | (-0.0384, 0.0413) |
| GAN200 | 0.70 | 0.24 | <0.01 | (0.1400, 0.3559) |
| GANLeast | 0.73 | 0.24 | <0.01 | (0.1081, 0.3058) |

The recall of GAN200 and GANLeast is under 0.75 and is significantly lower (p-value < 0.01) than the recall obtained for the non-enhanced virtual images. This drop in the recovery of the original anatomical structure indicates that enhanced images systematically distort the anatomical content. ContentNet recall is almost equal to the one obtained in virtual images with a difference less than 0.04, which shows that it consistently preserves the anatomical content along the sequence.

Concerning precision, ContentNet averages are a bit higher (p-value=0.025 and CI=(−0.1150, −0.0082)) than the average precision of original virtual images. This might be attributed to artifacts in the segmentation of distal airways that are prone to introduce shadows in virtual sequences resembling the appearance of luminal distal areas. Such shadowing artifacts do not appear in intra-operative videos and, thus, they are significantly reduced in ContentNet enhanced images. In fact, ContentNet precision ranges (0.94 ± 0.09) are closer to the full precision of intra-operative videos [12] than ranges for virtual images (0.8843 ± 0.1364). Finally, precision average ranges for GAN200 and GANLeast compares to virtual images precision which indicates that they present some kind of artifact (like the shadows of the images shown in Fig 4) that intra-operative videos do not have.

Fig 5 shows two representative examples of lumens detected on images enhanced with each method. Lumen centers are shown in blue points on, both, original virtual images and the enhanced ones. In the top case, GAN200 and GANLeast enhanced images miss the bottom branch. In the bottom case, in spite of a slight deviation in the location of its center, GANLeast succeeds in keeping the upper branch, while GAN200 enhancement has such a distorted anatomy that none of the original virtual branches can be identified. The proposed ContentNet preserves the virtual anatomy in both cases and, thus, all branches are properly detected.

**Fig 5.**
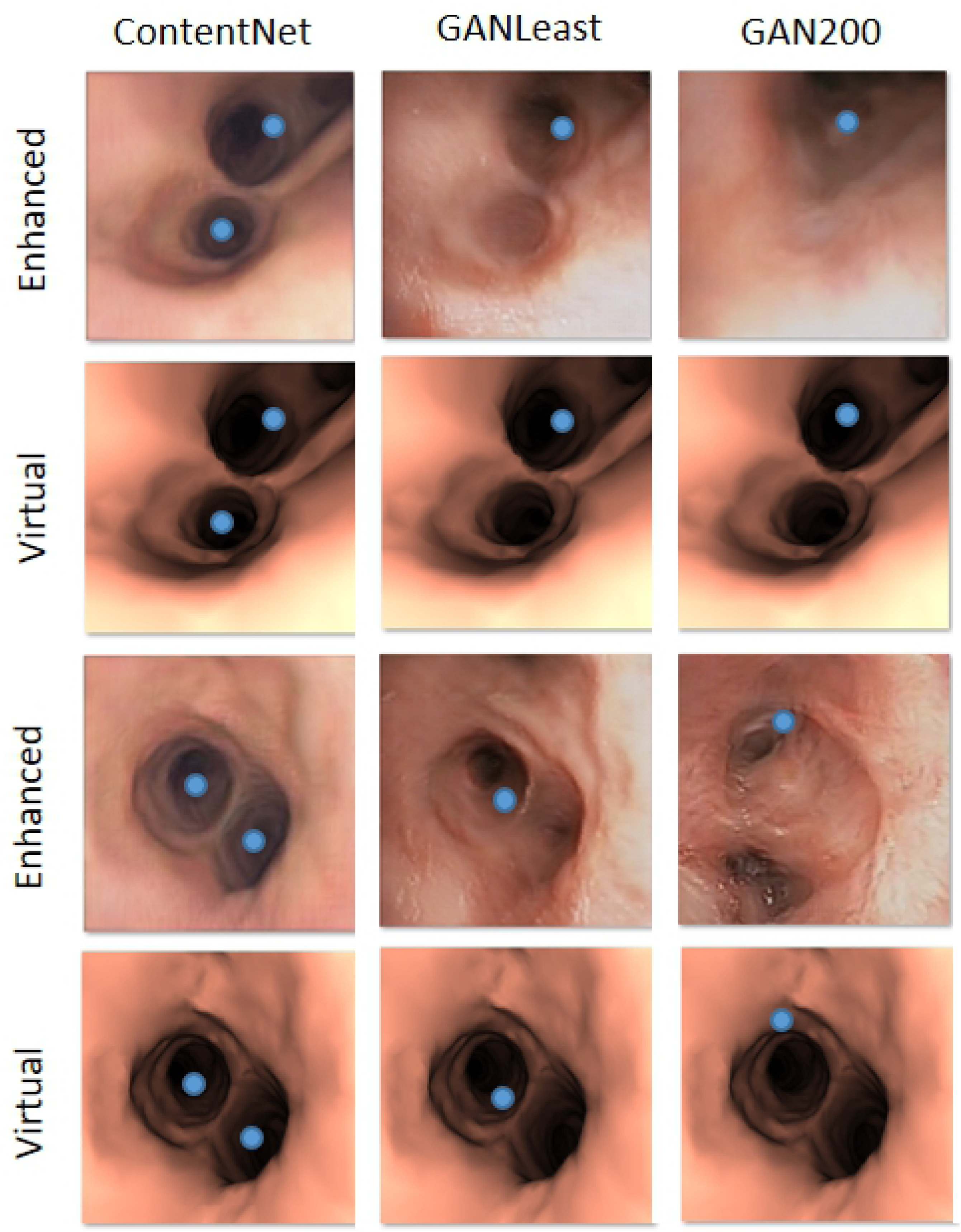
Detected lumens on virtual bronchoscopy enhanced using our method (ContentNet), 200th epoch network (GAN200) and the least cost one (GANLeast).

## Conclusion

We have presented a method for style transfer that preserves the structure of original input images and, thus, it is suitable for endowing virtual endoscopy with intra-operative appearance.

Content-net has been compared to state-of-art style transfer methods based on GANs. In particular, we chose cycleGAN after 200 epochs and the cycleGAN achieving the minimum cost. Two experiments were conducted. The first one assessed whether the appearance of enhanced images compares to intra-operative videos for their use in classification problems. The 2nd experiment assessed whether the anatomical content of the virtual images extracted from patient’s CT is preserved after their enhancement for their use in image processing problems.

Results obtained for the first experiment show that, like cycleGAN, Content-net has an appearance close enough to intra-operative videos as to be classified real by a discriminative network. This validates our method for data augmentation in classification problems. The 2nd experiment shows that images enhanced using both cycleGANs have a significant distortion in anatomical content (see Fig 5) and have larger temporal artifacts (see Fig 4) in comparison to Content-net. In fact, according to T-tests Content-net anatomical structure is not no significantly different from original virtual images extracted from patient’s CT anatomy. This validates our method for data augmentation in image processing problems.

The artifacts of cycleGANs images might be partially attributed to the adversarial training. On one hand, the combination of two loss functions with different (opposite, indeed) goals (minimization and maximization) introduces an oscillating behavior across training epochs and, thus, consecutive epochs might produce very different results. On the other hand, it is not guaranteed that both losses will have equal influence during training since the magnitude of one of the two might be predominant in the back-propagation of their gradients.

The proposed multi-objective approach allows the join optimization of both losses ensuring equal influence on the cycleGAN, regardless of their magnitude or gradient. This way, our multi-objective cycleGAN produces stylized images that share enough anatomical structure with virtual images as to be the input for a network blending both image pairs. The weighted skip connections of ContentNet provide selective blending of the structure and texture of these image pairs. This allows enhancing the patient specific anatomical content acquired by CT scans, while keeping an intra-operative appearance. In this context, it is worth noticing that ContentNet precision and recall ranges achieved in the detection of airways structure (centers) is very close to the ranges obtained in intra-operative videos [12].

In summary, two interesting conclusions can be inferred from our experiments. First, the use of multi-objective optimization strategies can be an effective alternative to back-propagation for the optimization of adversarial networks and other networks relying on multiple loss functions. In this context, the Pareto front condition could also be adapted for the selection of the most appropriate task in sequential multi-task learning. Second, the structure of neuron activations can be measured by the amount of information shared with input images. This measure of their content provides a description easy to interpret in terms of classical computer vision. We envision that this could be useful to define more specific and interpretable representation spaces based on CNNs.

